# A unique neurogenomic state emerges after aggressive confrontations in males of the fish *Betta splendens*

**DOI:** 10.1101/2020.08.05.237586

**Authors:** Trieu-Duc Vu, Yuki Iwasaki, Kenshiro Oshima, Masato Nikaido, Ming-Tzu Chiu, Norihiro Okada

## Abstract

Territorial defense involves frequent aggressive confrontations with competitors, but little is known about how brain-transcriptomic profiles change between individuals competing for territory establishment. Our previous study elucidated that brain-transcriptomic synchronization occurs in a pair-specific manner between two males of the fish *Betta splendens* during fighting, reflecting a mutual assessment process between them at the level of gene expression. Here we evaluated how the brain-transcriptomic profiles of opponents change immediately after shifting their social status (i.e., the winner/loser has emerged) and 30 min after this shift. We showed that unique and carryover hypotheses can be adapted to this system, in which changes in the expression of certain genes are unique to different fighting stages and in which the expression patterns of certain genes are transiently or persistently changed across all fighting stages. Interestingly, the specificity of the brain-transcriptomic synchronization of a pair during fighting was gradually lost after fighting ceased, because of the decrease in the variance in gene expression across all individuals, leading to the emergence of a basal neurogenomic state. Strikingly, this unique state was more basal than the state that existed in the before-fighting group and resulted in the reduced and consistent expression of genes across all individuals. In spite of the consistent and basal overall gene expression in each individual in this state, expression changes for genes related to metabolism, learning and memory, and autism still differentiated losers from winners. The fighting system using male *B. splendens* thus provides a promising platform for investigating neurogenomic states of aggression in vertebrates.

**Author summary:** Competitive interactions involve complex decision-making tasks that are shaped by mutual feedback between participants. When two animals interact, transcriptomes across their brains synchronize in a way that reflects how they assess and predict the other’s fighting ability and react to each other’s decisions. Here, we elucidated the gradual loss of brain-transcriptomic synchrony between interacting opponents after their interaction ceased, leading to the emergence of a basal neurogenomic state, in which the variations in gene expression were reduced to a minimum among all individuals. This basal neurogenomic state shares common characteristics with the hibernation state, which animals adopt to minimize their metabolic rates to cope with harsh environmental conditions. We demonstrated that this unique neurogenomic state, which is newly characterized in the present study, is composed of the expression of a unique set of genes, each of which was presumably minimally required for survival, providing a hypothesis that this state represents the smallest unit of neurogenomic activity for sustaining an active life.

## Introduction

Aggression is an evolutionarily conserved behavior and is used to establish and maintain a social hierarchy (1, 2). Many different animal species establish a social hierarchy to regulate social interactions among individuals competing for food, mating, and territory, which ensures the survival of the population (3). These hierarchical rankings lead to serious consequences for individuals including stress (4) and affect even the size of specific populations of nerve cells (5). Territorial defense involves frequent aggressive confrontations with competitors and has fitness consequences (6). Throughout the stages of territory establishment, individuals of different social status commonly express different sets of behaviors (a.k.a. behavioral states) that match their competitive ability. Thus, individual animals must be able to switch their social status, and this should be mediated by changes in gene expression in the brain that lead to distinct transcriptome profiles across the social behavior network (a.k.a. neurogenomic states) (7, 8). Previous studies have tried to map behavioral states onto neurogenomic states and have established these maps in various animals such as honeybees, songbirds, fish, and mice (9).

Fish—specifically trout, cichlid, and zebrafish—have recently become some of the most popular vertebrate models for dominance research because they possess easily observed dominance behaviors (e.g., chasing, biting, and gill flaring) (2), have recognition ability (10), and maintain their dominance hierarchies for extended periods of time (11). Despite the substantial genetic resources of these fish, studies using them have centered on dominance hierarchies and have explored how past experiences such as winning, losing, or observing a fight affect the outcome of their future encounters (11–15). Moreover, fighting interactions in these three species last for a short period of time, i.e., <36 min on average (8), and, therefore, the full range of volitional behavior expressed by one opponent toward the other for longer periods of time remains limited. Siamese fighting fish, *Betta splendens*, are notorious for their aggressiveness. Males of this species have stereotypical social displays that are well documented (16, 17), and individual fights can last for more than an hour. Studies using this fish have yielded several publications in the fields of ecology (18), pharmacology (19, 20), toxicology (21), metabolism (22, 23), and endocrinology including the study of sex hormones and steroid hormones (24, 25). Thus male *B. splendens* are suitable for investigating changes in neurogenomic states and behavioral states that occur across the stages of territory establishment.

The fighting process in *B. splendens* proceeds in a stereotypic manner, beginning with each fish displaying their fins and circling, followed by their performing a series of biting/striking/mouth-locking/chasing behaviors, and ending when one opponent shows fleeing/freezing behavior indicating surrender (26). Thus, one might expect that each fighting stage could have unique gene expression data associated with it (i.e., unique hypothesis) (27). Moreover, there are transient periods between these fighting stages, and some of the aggressive behaviors remain constant across stages, e.g., chasing. Thus, one might also expect to see that the brain transcription signal of a preceding fighting stage persists into subsequent stages, resulting in shared gene expression among fighting stages in the series. The expression of these genes would thus be either transient between temporally adjacent stages in the series or persistent across all fighting stages (i.e., carryover hypothesis) (27). Although these hypotheses help to improve our understanding of aggressive confrontation at the molecular level, we know little about which genes conform to a unique or carryover pattern across stages of fighting.

Here we have evaluated the unique and carryover hypotheses and have characterized the brain-transcriptomic responses of fighting opponents with a focus on the period after fighting concludes by mapping behavioral states onto neurogenomic states across stages of territory establishment. We experimentally stopped each fight at a certain fighting point, namely during fighting after 20 and 60 min had elapsed (D20 and D60, respectively) or immediately after and 30 min after fighting (A0 and A30, respectively) and analyzed brain transcriptome data from those fish, which allowed us to examine relationships between critical physiological events and behavioral decisions. A non-fighting group (B, before fighting) was also collected as a control group. Three analyses were performed: (i) brain-transcriptome changes across all fighting stages relative to non-fighting fish to capture the functional genes and molecular events accounting for changes in behavioral states; (ii) brain-transcriptome changes between individuals of interacting pairs to understand how fighting interactions were regulated across all stages; (iii) brain-transcriptome changes associated with winners and losers to obtain functional genes and molecular events underlying differences in neurophysiological changes between these two groups.

Moreover, the fighting interactions of paired male conspecifics are energetically costly, as indicated by an increase in the metabolic rate during the fighting process (22). Such energy consumption has a strong correlation with how they respond behaviorally, the strategy of which is evolutionarily conserved and has been observed across different animal species. For example, social challenge induces a rapid shift in energy neurometabolism and activates signals to drive neural plasticity in the brains of mice, honeybees, and stickleback fish (28). In the initial study, we had demonstrated that a vast number of genes are up-regulated in brains of *B. splendens* fish collected during fighting, showing a striking asymmetry in gene expression between the non-fighting and the fighting groups. These genes are associated with ion transport and metabolic processes, suggesting the possibility of a shift in metabolism that prefigures later neuronal plasticity in the brain (26). However, it remains unclear how energy depletion affects the changes that occur between transcriptomes of fighting opponents after their conflicts have resolved. Many mammalian species enter hypothermic states known as torpor or hibernation states to survive harsh environmental conditions such as food scarcity (29, 30). In this state, thermostatic animals actively reduce their basal metabolism to a minimum. We discuss here a possible link between the energy-saving survival strategy of fighting opponents after fighting and the hibernation state in thermostatic animals.

Here we showed that the unique and carryover hypotheses can be adapted to describe this fighting system and that a new and unique neurogenomic state emerges after fighting. Taken together, the results from our study lay out a comprehensive framework for investigating the neurogenomic states of aggression at the level of gene expression in vertebrates.

## Results and Discussion

### Behavioral dynamics of fighting

The stereotypical social displays during the first 60 min of a fight between male *B. splendens* were described in our previous study, in which each fighting stage was characterized by a particular set of behaviors (26) (Fig 1A). Briefly, fish performed display behaviors at the beginning of fights; during fighting, fish carried out biting, striking, surface-breathing, and mouth-locking; at the end of fighting, one fish chased the other; and 30 minutes after the chasing period, whereas the winners actively swam, the losers showed fleeing behavior indicating surrender. Here we measured the levels of behavioral repetition between opponents from seven fighting pairs and observed a significant difference between these levels enacted by the ultimate winners and losers across the antagonistic pairings. On average, the ultimate winners had a significantly higher number of bites/strikes relative to the ultimate losers during the initial 60 min (paired t-test: p = 0.02; Fig 1B), although total surface-breathing number did not significantly differ between the ultimate winners and the ultimate losers (paired t-test, p = 0.06; Fig 1C). However, there was a significantly positive correlation between biting/striking frequency and surface-breathing frequency (Pearson correlation test, r = 0.6, p = 0.03; Fig 1D).

**Figure 1.**
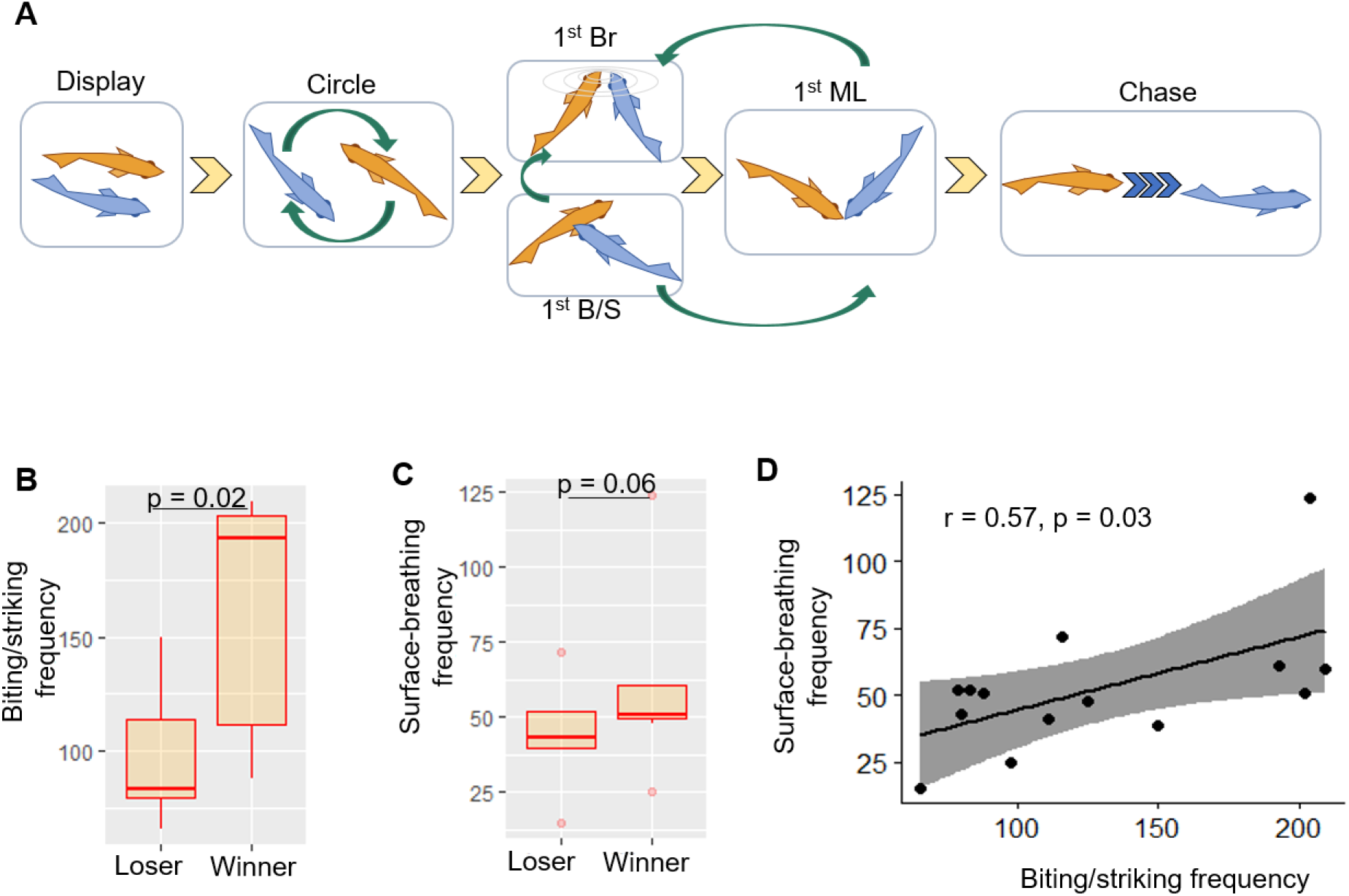
Behavioral dynamics of fighting. (**A**) Schematic illustration of dyadic fighting behaviors in male *B. splendens*. 1st Br, first surface-breathing; 1st B/S, first biting/striking; 1st ML, first mouth-locking. (**B** and **C**) Biting/striking frequency (B) and surface-breathing frequency (C) of winners and losers among male *B. splendens* fighting pairs given a 60-min fight duration. (**D**) Positive relationship between biting/striking frequency and surface-breathing frequency among male *B. splendens* fighting pairs given a 60-min fight duration.

The above observations agreed with several lines of evidence showing that more aggressive *B. splendens* fish tend to bite/strike more (8, 22, 31) and that two males of this species perform surface-breathing synchronously during interactions to meet oxygen requirements (16, 23). The synchronous surface-breathing between fighting opponents has also been observed in other air-breathing fish such as catfish (32) and pearl gourami fish (33). As low oxygen levels are typical habitat for *B. splendens* and for air-breathing fish (34), the surface-breathing behavior is likely an adaptation to cope with hypoxic conditions in these fish, and surface-breathing also results in a decrease in their gill surface area in comparison with other fish (35).

### Neurogenomic dynamics of fighting

The results obtained from behavioral analyses suggested the possibility of differences in brain signaling of the fighting opponents across the fighting stages. Therefore, we conducted RNA sequencing (RNA-seq) using whole brains collected at different fighting stages, namely B, D20, D60, A0, and A30, to observe the changes in gene expression. The principal component analysis using all genes (23,411 genes) across 37 brain samples showed that brain-transcriptomic responses of the fish were diverse during fighting but became highly consistent after fighting (Fig 2A). Then, we looked for differentially expressed genes (DEGs) by comparing all of the fighting groups (D20, D60, A0, A30) with the non-fighting group (B) (FDR < 0.05) and found that 6,758 genes (28.9% of the transcriptome) were differentially expressed as the sum of all of these comparisons (Fig 2B, S1 Table). In particular, the D60 and A0 groups showed more DEGs than the D20 and A30 groups, and the number of up-regulated genes was much higher than that of down-regulated genes within each fighting group (Fig 2B). In addition, the fold change (FC) in the expression level of each gene from a particular fighting group (D20, D60, A0, and A30) relative to that from the B group was calculated for the DEGs. Their distributions are shown in Fig 2C, in which the median FC value was significantly different across all the fighting groups (ANOVA, p < 22e-16) (S2 Table), suggesting that each fighting group has a particular set of transcriptomic characteristics.

**Figure 2.**
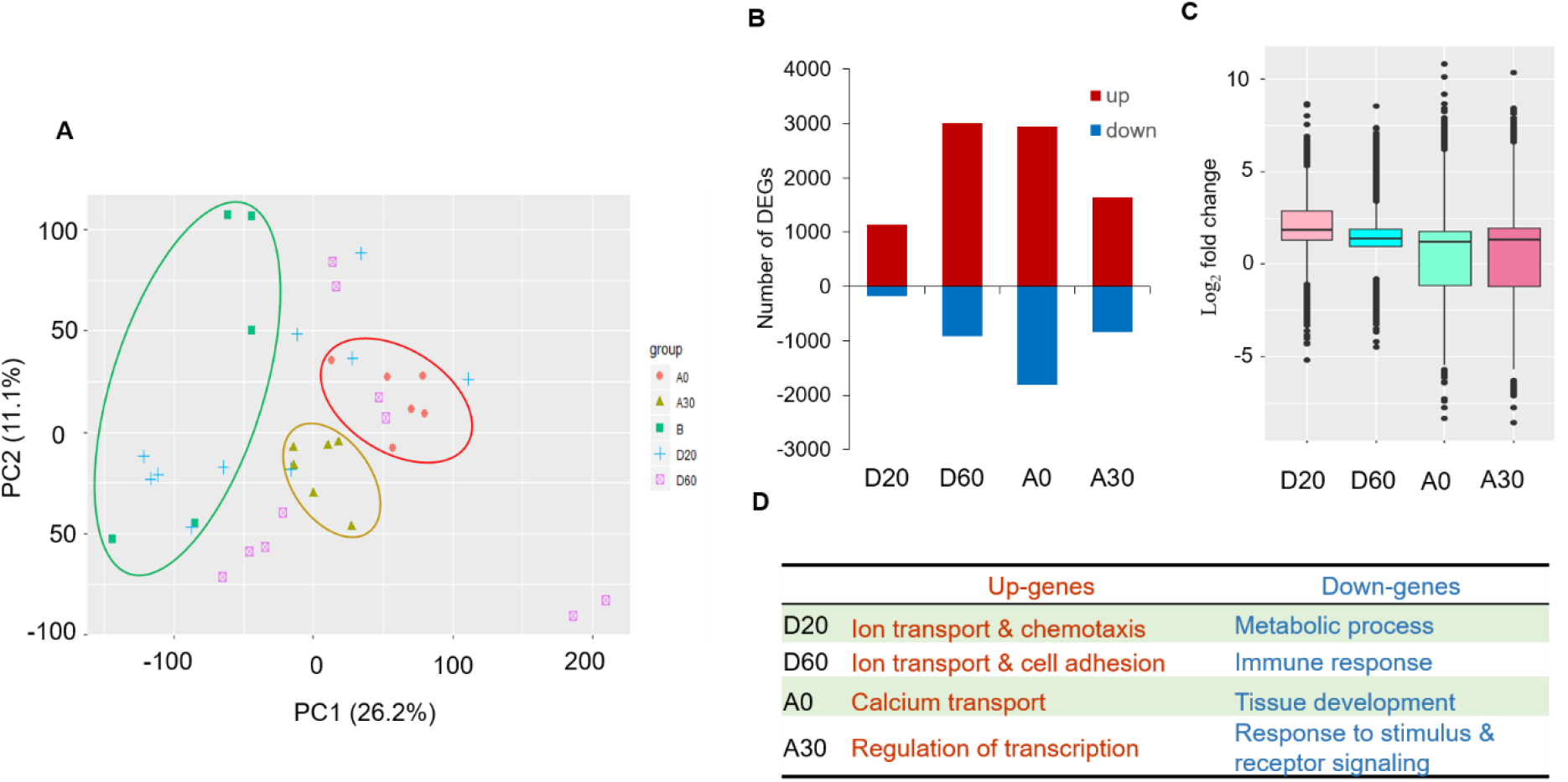
Neurogenomic dynamics of fighting. (**A**) Principal component analysis (PCA) using all 23,411 genes across 37 samples. The circled data points indicate transcriptomes from a fighting group that was more stable. (**B**) The number of up- and down-regulated differentially expressed genes (DEGs) of each fighting group relative to the non-fighting group. (**C**) A boxplot showing the relative fold change (FC) of DEGs for each fighting group relative to their expression in the non-fighting group. (**D**) Summary of gene ontology (GO) terms that were enriched in up-regulated genes (up-genes) and down-regulated genes (down-genes) of each fighting group.

Next, we conducted functional enrichment analysis for subsets of the above DEGs with more stringent criteria (FDR < 0.05 and |logFC| > 2) using DAVID and REVIGO. Broadly, fighting coincided with changes in ion transport in the brain along with modifications of the immune response and transcription. Genes associated with metabolism and the immune response were down-regulated in the D20 and D60 groups relative to the B group. Genes associated with calcium signaling were up-regulated in the A0 group as the fish shifted their social status (one chasing the other). Genes associated with transcription regulation were up-regulated in the A30 group, and genes associated with ion transport were up-regulated in all fighting groups except the A30 group (Fig 2D, S3 Table).

Last, we characterized the expression of immediate early genes (IEGs) because their expression reflects the recent activity of neurons (36). We manually screened references with the annotation data associated with the DEGs and obtained a list of 44 IEGs whose expression patterns were classified into three groups—group I (highly expressed in D20), group II (highly expressed in D60), and group III (highly expressed in both D20 and D60)—across the B, D20, and D60 fish (Fig 3A). Then, we further investigated the expression of *c-fos*, which belonged to the group I IEGs and is one of the most representative molecular markers of neural activity (37). We observed that the relative *c-fos* expression levels showed significant differences across the different fighting stages, except for the A0 vs. A30 and D60 vs. A30 fish, and returned nearly to their non-fighting levels once the fight ended (ANOVA, p < 0.05) (Fig 3B). Subsequently, in situ hybridization using a *c-fos* probe was performed to examine neuronal activity in neurons of the D20 fish. Expression of *c-fos* was detected in the forebrain of the D20 fish (Fig 3C), which contains loci known to play key roles in the control of the fear response, spatial learning, and reproduction (38), suggesting that these fish experienced fear as well as anxiety and showed cognitive ability during fighting. Collectively, the above observations suggest that the fighting involved significant neurogenomic shifts in *B. splendens* males.

**Figure 3.**
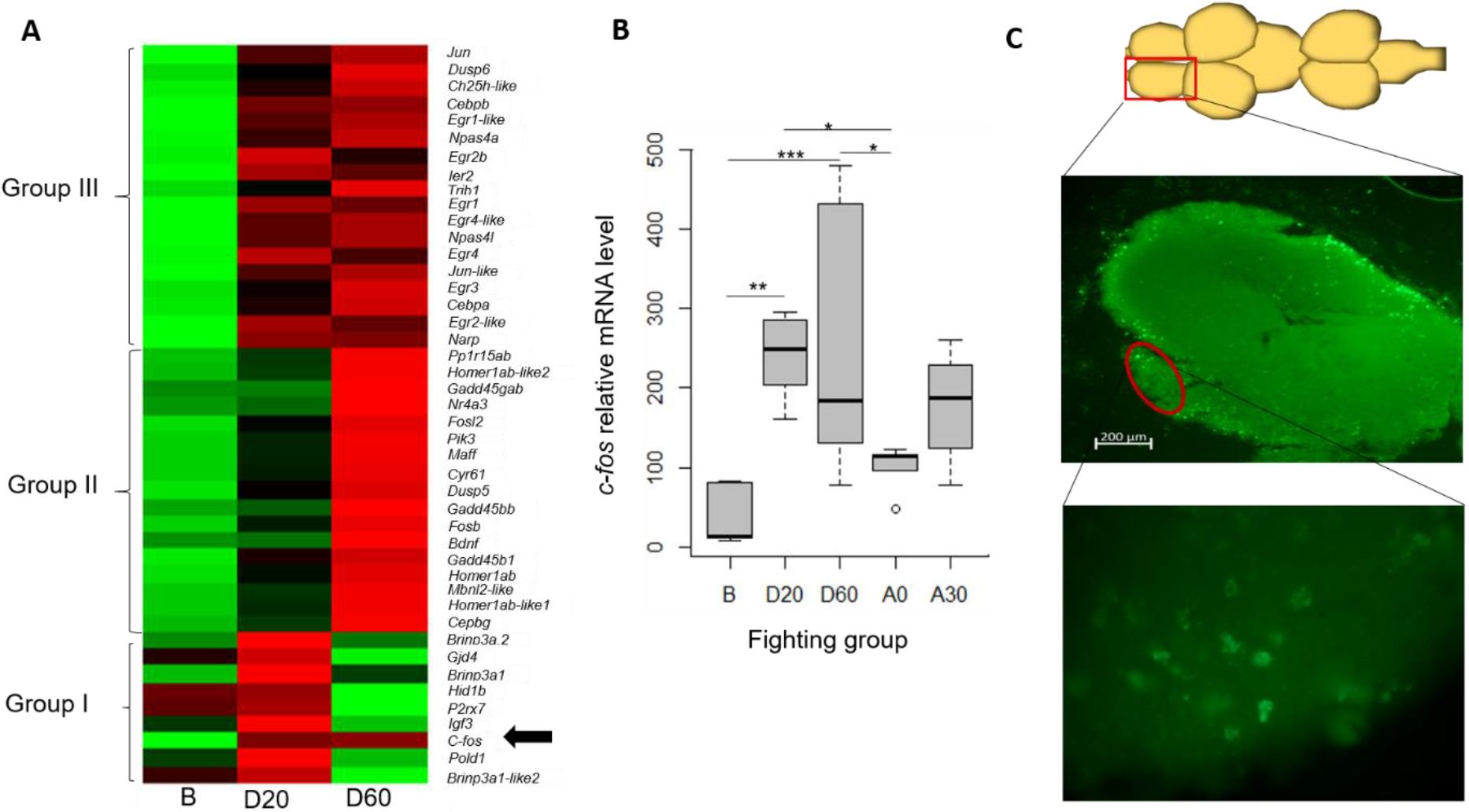
*c-fos* expression as an indicator of neuron activity. (**A**) Heatmap depiction of expression profiles of the immediate early genes (IEGs), which were divided into three groups as described in the Results. Color intensity indicates the level of expression (red, high expression; green, low expression); the *c-fos* gene is marked by an arrow. (**B**) Relative expression of *c-fos* gene across all fighting groups. Statistical significance between groups was assessed by an ANOVA followed by a post hoc Tukey’s test. *p < 0.05, **p < 0.01, ***p < 0.001. (**C**) Mapping of *c-fos* expression based on in situ hybridization. The upper image illustrates the entire fish brain (dorsal view, with anterior to the left). The middle image shows a brain section taken from the forebrain region of a fish from the D20 group, with a higher-magnification image of the indicated region in the lower image.

### Changes in and stability of the neurogenomic state across fighting stages

Fig 1A showed that each fighting stage is characterized by a particular set of behaviors, and so we hypothesized that each fighting stage might have a unique neurogenomic state associated with it (referred to as the unique hypothesis). To test this possibility, we first looked for whether certain DEGs were specific to each fighting stage, i.e., not shared with other fighting stages (see Methods). To obtain this, we generated a Venn diagram by overlapping the four gene lists of the above DEGs obtained from four fighting groups against the non-fighting group (FDR < 0.05, Table 1).

**Table 1.** Overlapping genes between fighting stages

| Comparison<br>(group1 vs.<br>group 2) | Overlapping genes | Group 1 total DEGs | Group 2 total DEGs | p-value |
| --- | --- | --- | --- | --- |
| D20 vs. D60 | 922 | 1,318 | 3,912 | 0 |
| D20 vs. A0 | 966 | 1,318 | 4,756 | 0 |
| D60 vs. A0 | 2,718 | 3,912 | 4,756 | 0 |
| D20 vs. A30 | 739 | 1,318 | 2,480 | 0 |
| A0 vs. A30 | 1,806 | 4,756 | 2,480 | 0 |
| D60 vs. A30 | 1,612 | 3,912 | 2,480 | 0 |

Each overlapping gene set was assessed and its corresponding p-value was generated by the hypergeometric test

Consistent with the unique hypothesis, this analysis resulted in—with high statistical confidence— lists of 184, 891, 1,528, and 411 DEGs that were uniquely expressed in the D20, D60, A0, and A30 fish, respectively (Fig 4A). The expression patterns of these uniquely expressed DEGs in each fighting stage are shown in Fig 4B, clearly confirming their stage-specific expression. Interestingly, we noticed that the considerable number of down-regulated genes (289 genes) in the DEGs unique to A0 (among a total of 1,528 genes) were in sharp contrast to their up-regulated expression in the D20 fish (Fig 4B, the third column). We thus referred to these 289 unique DEGs as “geared fighting genes” given that they were highly expressed in the D20 group to put these fish in gear for fighting, but they were shifted to neutral in the A0 group, as the fighting interactions had been resolved (S4 Fig, S4 Table). These geared fighting genes were associated with pigmentation (*tjp1a*, *atoh7*, and *tyrp1b*), proteolysis (*mmp16b*, *usp43a*, *dpp6b*, and others), and striated muscle cell differentiation (*six1b* and *pobdc2*). Regarding the coloration changes, our initial analysis showed that both fighting opponents spent a considerable proportion of the initial 20 min of the fight (~70%) in changing their color patterns i.e., switching to a darker or lighter shade (S5 Table). Whereas coloration has important roles in the immune response, sexual discrimination, and fighting coordination in *B. splendens* (39), sticklebacks (40), and cichlids (41) at the behavioral level, the involvement of pigmentation-related genes in this study suggested the molecular basis for color signaling in territorial aggression.

**Figure 4.**
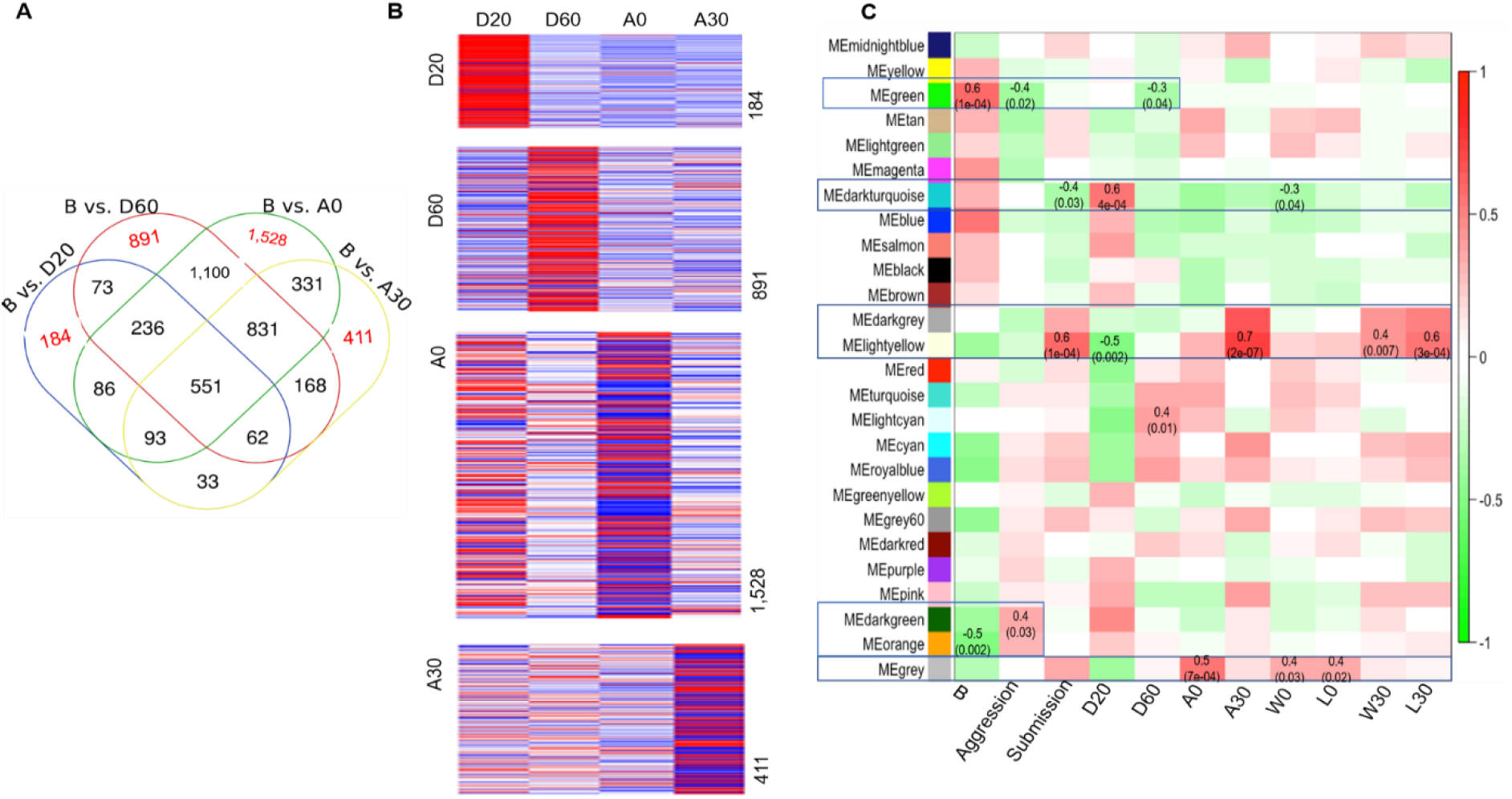
Changes in and stability of the neurogenomic state across stages of fighting. (**A**) Venn diagram showing the overlapping DEGs between each fighting group (D20, D60, A0, A30) relative to B, the non-fighting group. The unique gene sets of each fighting group were highlighted in red (**B**) Heatmap depiction of expression profiles of the genes that were differentially expressed in only one fighting stage. Color intensity indicates the level of expression (red, high expression; blue, low expression). (**C**) Associations between patterns of expression in 26 identified modules across all fighting groups (B, D20, D60, A0, and A30) and winners/losers (W0, L0, W30, and L30) and across the observed behavioral traits aggression (biting/striking/chasing) and submission (fleeing**/**freezing). Each row corresponds to a module eigengene (ME), and colors represent the correlation coefficient between the ME and fighting groups as well as between the ME and behavioral traits. The colors of the boxes are scaled with the value of correlation coefficients, ranging from green (r = −1) to red (r = 1). Some of the values for r (top number) and p-values (bottom number in parentheses) are shown. The values of r represent the degree of correlation, in which those values with a negative integer indicate a negative correlation and those values with a positive integer indicate a positive correlation. The five blue boxes indicate that these ME values are the most significantly associated with the social phenotypes as well as the behavioral traits.

Additionally, all gene sets identified across all fighting stages (23,411 genes) were further analyzed with a weighted gene co-expression network analysis (WGCNA) to identify the gene modules of co-expressed genes associated with each fighting group (B, D20, D60, A0, and A30). This approach identified 26 distinct gene modules, and significant correlations were found for 9, 11, 4, 5, and 5 modules in the B, D20, D60, A0, and A30 groups, respectively (Fig 4C; S6 Table). Together, the DEGs and WGCNA results suggest that the unique hypothesis can be applied to this system in which certain genes were DEGs only within a particular stage. However, whether their differential expression results in a subsequent behavior or is a consequence of a behavior that had already happened remains unknown.

Next, we assessed the extent to which genes were shared among different fighting stages by testing whether the possibility of overlapping DEGs between fighting stages (Fig 4A) was greater than expected by chance using the hypergeometric test. Consistent with the carryover hypothesis, the number of overlapping DEGs between fighting stages was statistically different across fighting stages, and they showed substantial correlation in expression across these six comparisons (Table 1, S5 Fig). Those shared DEGs between adjacent fighting stages, i.e., D20 and D60, D60 and A0, and A0 and A30, were referred to as transitional genes, which might be involved in facilitating the transition into the next fighting stage, priming for and/or responding to a particular event or stimulus during that fighting stage, e.g., a shift in social status. In addition, we used a more stringent criterion than was used in the analysis of Fig 4A (FDR < 0.05 and |log FC| > 2 instead of only FDR < 0.05) to look for the overlapping genes and found 227 up-regulated genes as well as seven down-regulated genes, each of which was expressed in the same direction, i.e., up- or down-regulated, across all fighting stages (S6A Fig and S6B Fig). These genes were referred to as persistent genes, and they may be involved in maintaining the previous neurogenomic state and/or reflect the constant demands of fighting, e.g., chasing must be maintained across all stages of fighting. Similar unique, transient, and persistent gene expression patterns have been previously observed in studies of the response to an intruder or of parental care in sticklebacks (27, 42), suggesting that some of the molecular mechanisms of aggressive behavior might be deeply conserved.

### A unique neurogenomic state emerges after fighting

Our previous study revealed that the brain-transcriptomic patterns of fighting opponents become synchronized during fighting (i.e., among the D20 and D60 fish) (26). Here, we asked how these brain-transcriptomic patterns changed after conflicts were resolved (i.e., among the A0 and A30 fish). Similar to our previous analysis among the ‘during’ fighting groups (26), we calculated and compared the correlation coefficients of expressed genes (r values) between paired fish and unpaired fish for the A0 group (i.e., A0 paired fish vs. A0 unpaired fish) and for the A30 group (i.e., A30 paired fish vs. A30 unpaired fish) (see Methods). The r values for these two comparisons were not significantly different and remained high for both paired and un-paired fish in both groups, whereas those between paired fish and unpaired fish during fighting were significantly different (especially in the case of D60), as was described previously (26) (Fig 5A). Interestingly, based on a clustering analysis using all 23,411 genes presented in a heatmap, the paired fish of A0 were clustered in a pair-specific manner, as was the case for D60 (26). In the case of A30, however, both paired- and unpaired fish were grouped into a single cluster (Fig 5B) and showed no pair-specific characteristics. Furthermore, the grade of membership (GoM) results indicated that the pair-specific synchronization pattern, which was still observed in the A0 fish to some extent, was completely lost in the A30 fish (Fig 5C; W0 and L0 data vs. W30 and L30 data). In fact, the ratios of each component in the GoM among the six A30 individuals, including paired and unpaired fish, looked very similar. Noticeably, the third component (orange color) in the GoM analysis occupied the largest proportion among the seven components in the A30 fish. The top 100 driving genes of this component (i.e., the genes that were most distinctively differentially expressed) were associated with membrane transport and voltage-gated potassium channel activity (S7 Table). It is likely that a neurometabolic shift was accompanied by changes in potassium channel expression. This process may play a role in coping with stress responses as has been previously reported in mice (43).

**Figure 5.**
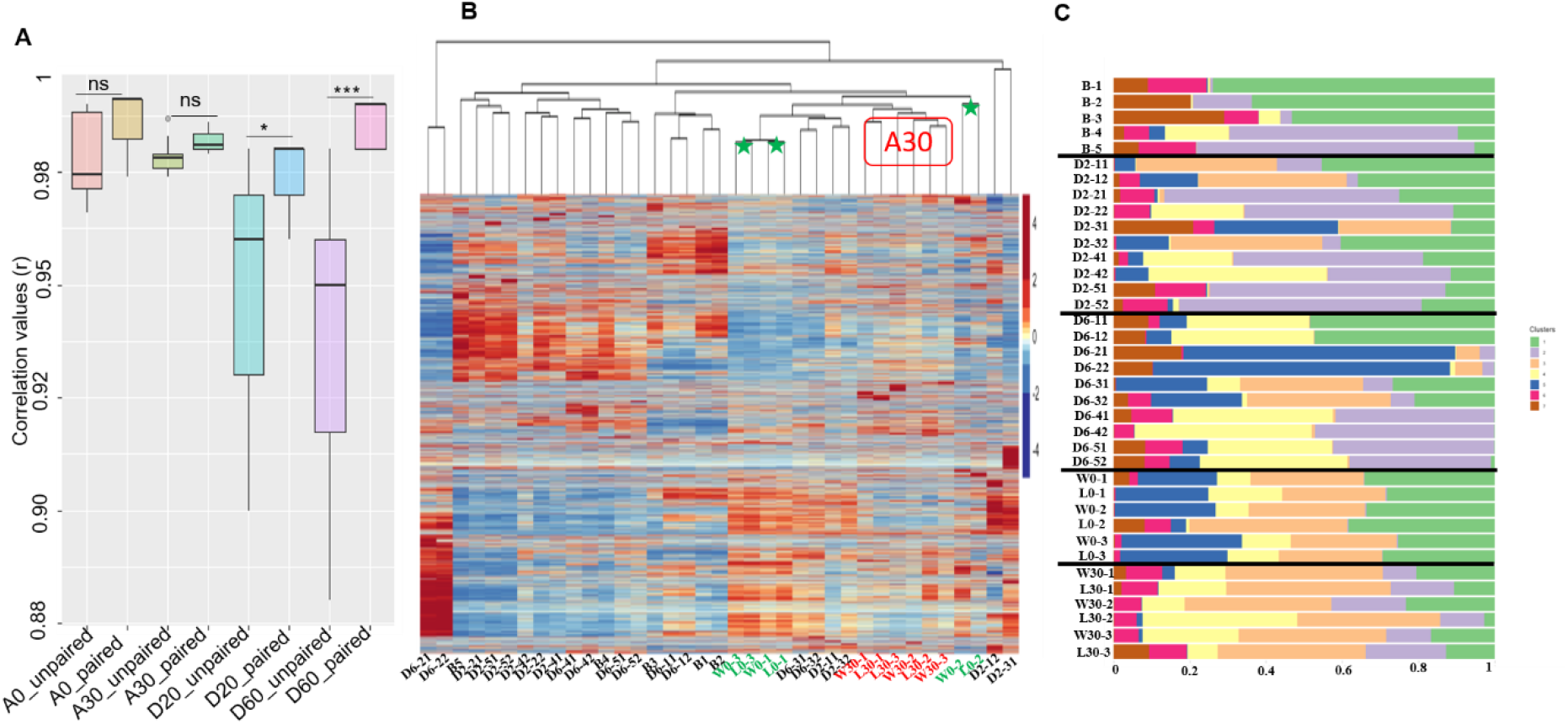
A unique neurogenomic state emerges after fighting. (**A**) Correlation of gene expression values between paired fish vs. unpaired fish among all fighting stages. ns, not significant, *p < 0.05, ***p < 0.001. (**B**) Heatmap using the 37 cDNA libraries with each of the 23,411 gene contigs. The color intensity indicates expression level (red, high expression; blue, low expression). All sample names can be seen at the bottom of the figure. Samples of the A30 fighting group are marked in red. Two paired A0 fish are marked with green stars. (**C**) GoM analysis using TMM values for all 23,411 genes generated from the 37 brain samples with a set of seven clusters represented by different colors.

Then, we examined how the expression of each gene varied among individuals within each fighting group. We hypothesized that the different fighting experience of each opponent in a fighting pair may be reflected in the variable expression of certain genes. As a measure of variance, we used the coefficient of variation (CV), which was computed for each gene (44). We showed that the A30 fish had the smallest CVs, whereas the D60 fish had the largest CVs, and the CV values were significantly different for all comparisons between two groups including the fighting as well as the non-fighting fish, except for the comparison between D20 and D60 (ANOVA, Fig 6A, S8 Table). Strikingly, an analysis of the expression patterns of the 15 most variable genes across all 37 individuals showed that their expression showed the least variability in the A30 fish (Fig 6B), consistent with our earlier observations based on the heatmap (Fig 5B) and GoM (Fig 5C) analyses. These genes include *neutrophil elastase* (*ela2*), which plays a role in degenerative and inflammatory diseases (45); *glutathione S-transferase omega* (*gsto1*), which is associated with the stress response (46); and *intersectin 1* (*itsn1*), which is involved in synaptic vesicle recycling (47), among others.

**Figure 6.**
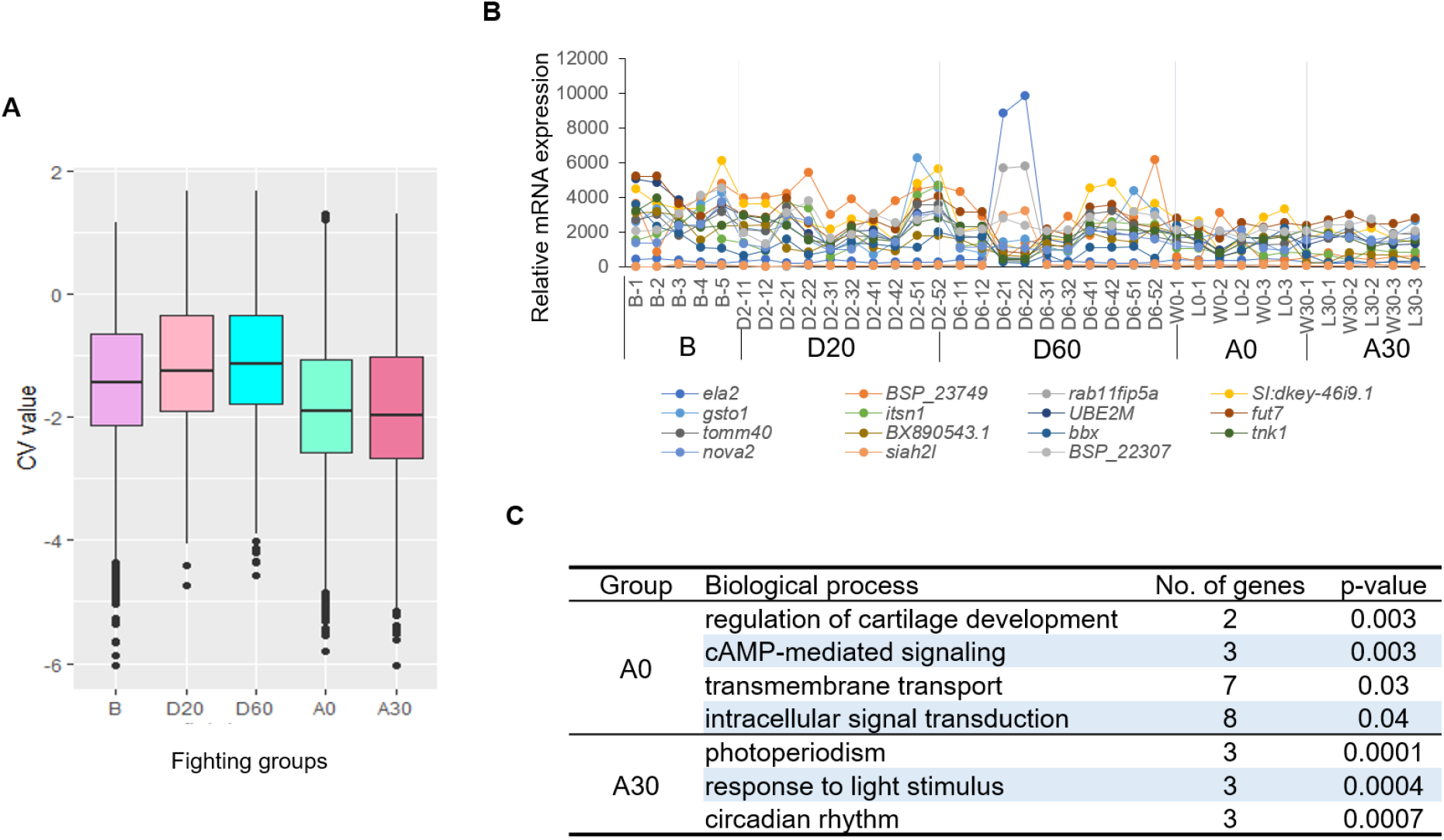
Fighting experience results in different expression variance in brain transcriptomes. (**A**) Coefficients of variation (CVs) for individual genes from the whole-brain transcriptomes (n = 23,411 genes) for B, D20, D60, A0, and A30 groups. (**B**) Expression patterns of the 10 most variable transcripts across all 37 samples. (**C**) Enriched biological process terms for the synchronized genes found in the A0 and A30 fish.

Next, we identified genes that were synchronized between individuals of the paired fish in the A0 and A30 groups. We obtained 244 synchronized genes (associated with signal transduction, cAMP-mediated signaling, and transmembrane transport) for the paired A0 fish and only 101 synchronized genes (associated with circadian rhythm, response to light stimulus, and photoperiodism) for the paired A30 fish (Fig 6C, S9 Table). The number of synchronized genes between the paired fish after fighting was much smaller than those between the paired fish during fighting, i.e., 1,522 genes for the D60 group (26). The loss of pair-specific synchronization of brain transcriptomes in the A30 fish could be due to the absence of fighting interactions between opponents after the fight was concluded. This loss of pair-specific synchronization, as well as the minimal variability in gene expression in the A30 fish (Fig 6B), the clustering by heatmap (Fig 5B), and the composition similarity based on GoM analysis (Fig 5C), provides the basis for our proposal of the emergence of a new and basal neurogenomic state in the A30 fish, in which the expression level of each gene was suppressed to the necessary minimum level due to exhaustion after the fight. As it would be expected to consider that the behavioral state of the before fighting group corresponds to a basal neurogenomic state, the discovery of this new and more basal neurogenomic state in our study is really surprising and requires further investigation.

In the case of the basal neurogenomic state in the A30 fish, the expression of each gene was presumed to be suppressed to the necessary minimum level due to exhaustion after the fight, as discussed above. We propose here that this state appears to be similar to a torpor (or hibernation) state, in which thermostatic animals actively reduce their basal metabolism to a minimum to survive harsh environmental conditions, e.g., food scarcity (29, 30). Recently, researchers demonstrated that the torpor state in rodents is regulated by neurons in the hypothalamus that express pyroglutamylated RF-amine peptide (30). This neuronal population is located in the anteroventral periventricular nucleus (AVPe) and medial preoptic area (MPA) of the rodent brain (30). Other researchers characterized 39 torpor-regulated genes in mice based on single-cell sequencing (29). Interestingly, we found nine of these torpor-regulated genes that overlapped with the DEGs of the A0 group relative to non-fighting group, among which five genes—*pde10a* (associated with signal transduction)*, adgrb3* (associated with the formation of new blood vessels)*, fndc3a* (associated with epidermal cell regeneration)*, adarb2* (RNA-editing enzyme), and *eprs* (charged-tRNA enzyme)—are specific to the A0 group (S10 Table). Future experiments to examine the temporal shift in the expression of these genes during fighting in the AVPe and MPA brain regions using in situ hybridization or using an RNA-seq approach specifically with the analogous regions in the fish brain (instead of using the whole brain) may help us to understand the molecular mechanism**s** responsible for the emergence of the basal neurogenomic state in *B. splendens* after fighting. Furthermore, understanding the roles of these genes in the torpor state might suggest an evolutionary process by which the thermostatic system of mammals developed from other pathways existing in non-thermostatic animals such as fish.

### Genes and molecular events associated with the emergence of winners and losers

Based on the WGCNA results (Fig 4C), both W0 and L0 were characterized by the grey module (positively associated), and W0 was also characterized by the darkturquoise module (negatively associated). In contrast, both W30 and L30 were characterized by the darkgrey module (positively associated) and lightyellow module (positively associated) (Fig 4C, S6 Table). Genes in these modules were associated with different biological process terms (S7 Fig). Moreover, aggressive behaviors (biting/striking/chasing) were negatively associated with two modules (green and tan) and positively associated with two modules (darkgreen and orange), and submissive behaviors (freezing/fleeing) were positively associated with three modules (grey, darkgrey, and lightyellow) and negatively associated with one module (darkturquoise) (S6 Table). Together, these results suggest the involvement of different transcriptional networks in the observed phenotypes for winners and losers.

Next, we compared gene expression in A0 and A30 fish relative to that in D60 fish (FDR < 0.05 and |logFC| > 2), given that critical neurophysiological events and behavior decisions may be initiated at D60. The number of down-regulated genes was much higher than that of up-regulated genes in the after-fighting groups (W0, L0, W30, and L30) relative to the D60 group, and there were very few DEGs between winners and losers at the two time points after fighting (Fig 7A). A KEGG pathway analysis for these DEGs showed that both winners and losers (W0, L0, W30, and L30) were enriched for metabolic-related genes, i.e., *ckmb*, *pygma*, and *pkma* kinases, which are involved in generating ATP and in the response to hypoxia tolerance in the brain (48). This suggests that the potential for metabolic activities and the hypoxia tolerance of both opponents decreased after fighting. Among down-regulated genes in the four groups examined after fighting, only those in the L30 group were associated with calcium signaling, the MAPK pathway, and cell adhesion molecules (Fig 7B), all of which are associated with learning and long-term memory (49). Furthermore, we performed gene enrichment analysis for the L30 and W30 DEGs and showed that L30 enriched genes were related to cell adhesion, signaling regulation, and ion transport, whereas W30 enriched genes were related to pre-synaptic organization and neuronal cell-cell adhesion (Fig 7C). Collectively, these findings suggest a close link between a shift in neurometabolism and behavioral plasticity, which is supported by evidence that social challenge induces a rapid shift in neurometabolism in the mouse leading to neural plasticity in brains of mice (43) and in honeybees (50).

**Figure 7.**
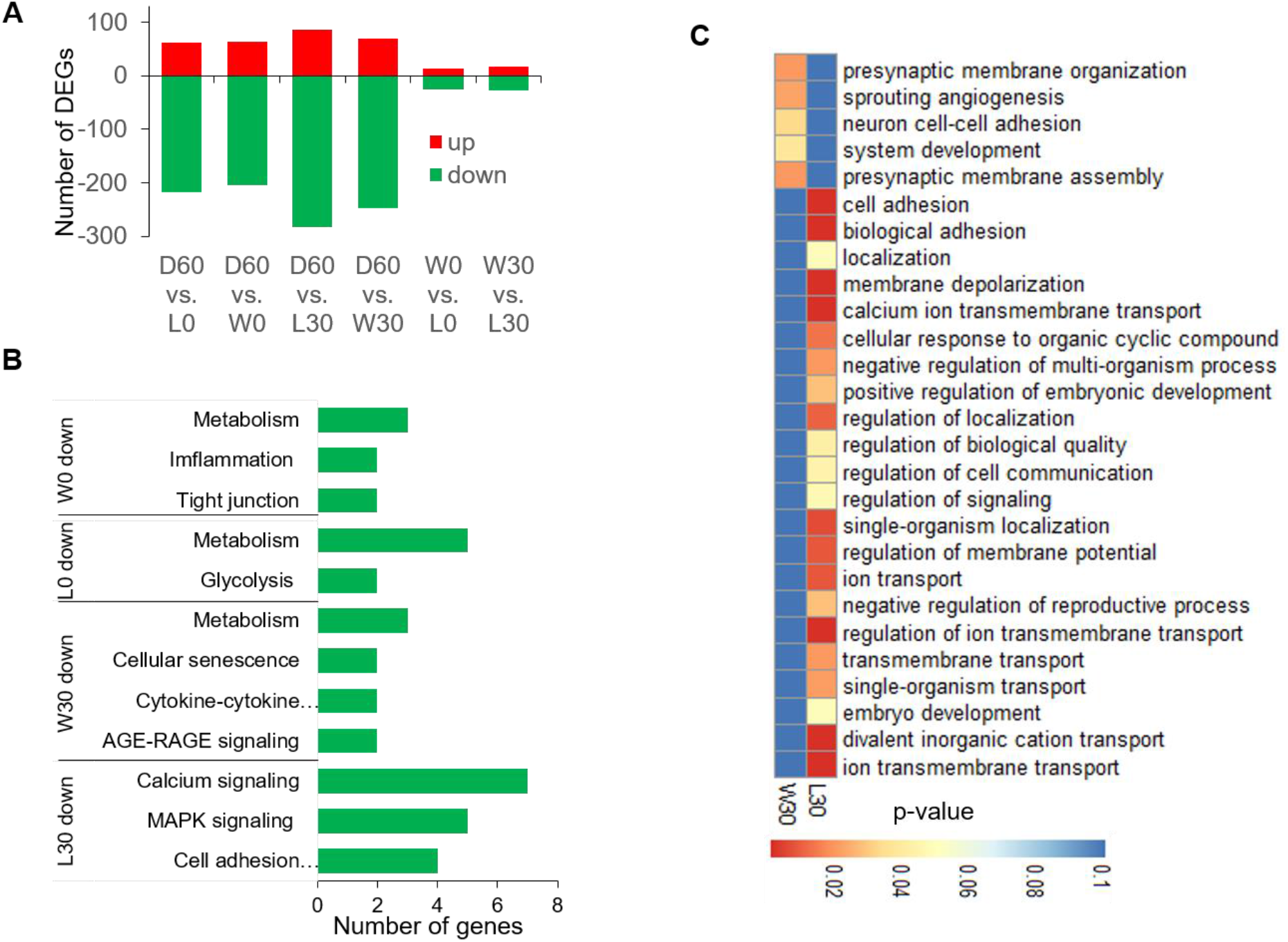
Shift in social status and corresponding changes in gene expression. (**A**) Comparison of DEGs generated from the after-fighting groups (W0, L0, W30, and L30) relative to D60 and between winners and losers (W0 vs. L0 and W30 vs. L30). Red refers to up-regulated genes, and green refers to down-regulated genes in the A0 and A30 fish relative to D60 fish. (**B**) KEGG pathway enrichment analysis for the DEGs obtained from the W0 vs. D60, L0 vs. D60, L30 vs. D60, and W30 vs. D60 comparisons. (**C**) Enriched biological process terms for the DEGs obtained from the D60 vs. L30 and D60 vs. W30 comparisons.

### Genes and molecular events associated with neurophysiological differences between winners and losers

The fish *B. splendens* has been shown to have cognitive ability based on the observation that they are able to gather information after viewing contests between neighbors and can use this information for future fights (10). Thus, we determined whether fighting opponents were able to encode their previous fighting experience. We thus searched for learning- and memory-associated genes among the DEGs of the W0, L0, W30, and L30 fish relative to the B fish by using the UniProt database—a comprehensive resource for protein sequence and annotation data. Losers showed a greater enrichment for learning and memory genes than winners, i.e., 357 for L0 vs. 64 for W0 and 276 for L30 vs. 176 for W30 (FDR < 0.05; S11 Table). Overlap between these gene lists revealed a greater number of loser-specific genes than winner-specific genes, i.e., 183 L0-specific genes vs. 3 W0-specific genes and 78 L30-specific gene vs. 24 W30-specific genes; there were 29 genes that were found in all four groups of fish (S8 Fig). A KEGG analysis of these specific genes showed that only the L30 genes were enriched for oxidative phosphorylation and ErbB pathways, which are associated with learning and memory (51); no KEGG pathways were found for the W0-, L0-, or W30-specific genes. Collectively, these results suggest that genes associated with learning and memory exert more influence on losers than winners. The involvement of these genes (S9 Fig) could help fighting opponents to encode their fighting experience and to use that information to guide future behavior (43, 52).

Further, we noticed that at 30 minutes after the end of fighting, while the winner (W30) was actively swimming, the loser (L30) showed reduced interactions, performed repetitive behaviors, and showed a higher level of inertness (Video S1). As these behavioral displays overlapped with typical behaviors among human individuals with autism spectrum disorder (ASD) (53), we investigated whether there are any genes in the W30 or L30 fish that are orthologous to ASD-risk genes in humans. We first obtained the lists of DEGs from the B vs. W30 and B vs. L30 comparisons, as well as the list of ASD-risk genes from the SFARI database (54). Then, we looked for overlap between the DEGs and the ASD-risk genes and detected a total of 35 ASD-risk genes that are also DEGs. Among these 35 ASD-risk genes, 19 genes had orthologs among the W30 DEGs, and 28 ASD-risk genes had orthologs among the L30 DEGs. By looking for overlap with these 19 and 28 ASD-risk genes, we found 7 W30-specific ASD-risk genes, 16 L30-specific ASD-risk genes, and 12 ASD-risk genes that were DEGs in both W30 and L30 fish. Whereas most of the L30-specific ASD-risk genes (10/16 genes) were categorized as having high and strong evidence (score 1 and 2), all of the W30-specific ASD-risk genes were listed as suggestive evidence (score 3 and S) (Fig 8A). The expression patterns of some ASD-risk genes that were specific for L30 and W30 are shown in Fig 8B and Fig 8C, respectively. Several of these ASD-risk genes should be noted when considering the etiology of ASD. For example, *cacna1c* mediates the influx of calcium ions into the cell upon membrane polarization (55), *ctnna2* regulates the morphological plasticity of synapses during development (56), and *cdh13* is a negative regulator of axon growth during neural differentiation (57). These ASD-risk genes have been associated with autistic-like behavior in honeybees (58), cave fish (59), and mammals (58). Together, the potential for shared genetic underpinnings between losers and individuals with ASD could provide further insights into the cause of ASD (60) and may shed light on the influence of social interactions on ASD.

**Figure 8.**
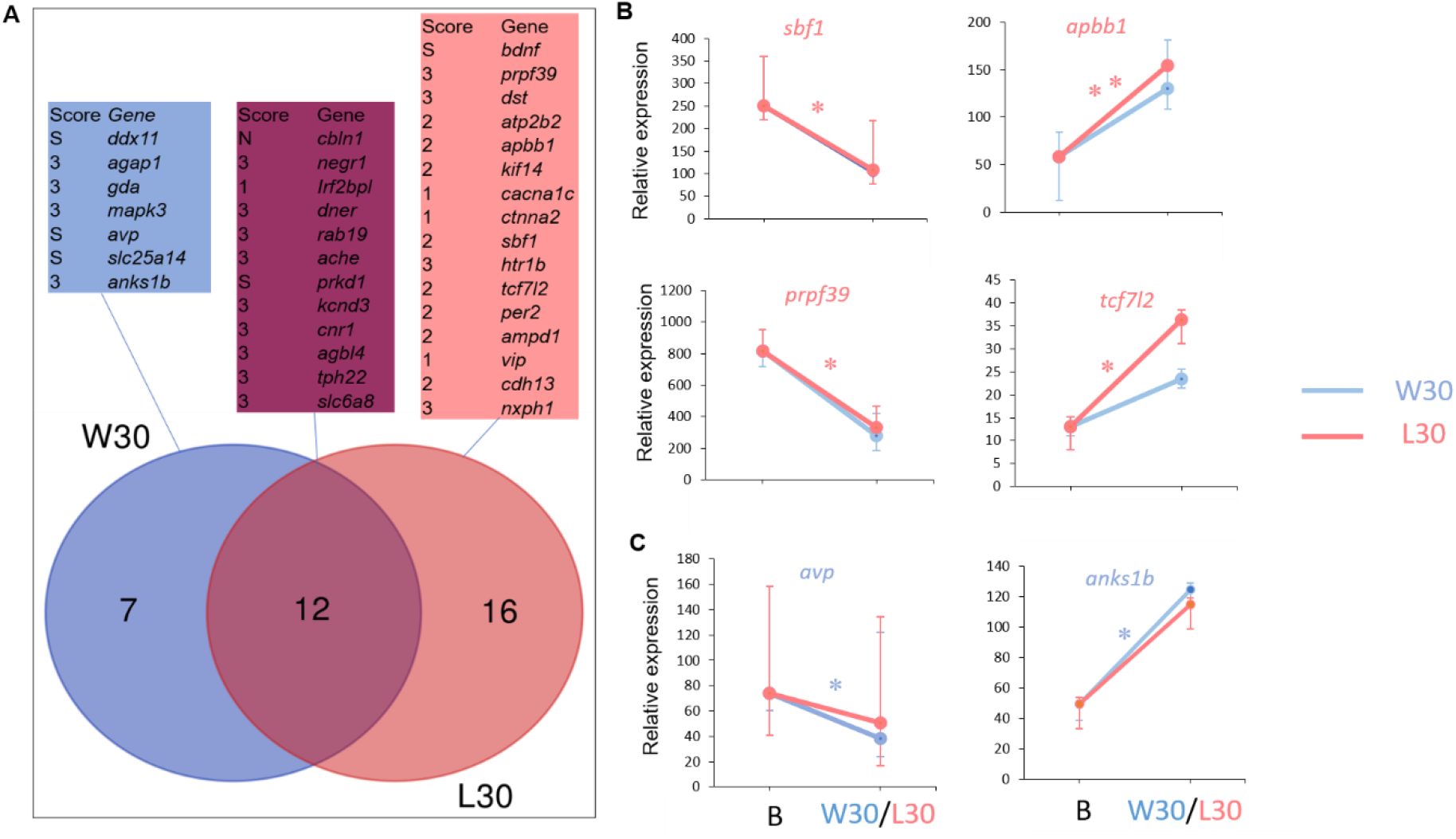
Autism risk genes found in W30 and L30 DEGs. (**A**) Venn diagram showing the number of genes associated with increased risk of autism found among W30 and L30 DEGs relative to the non-fighting fish. ASD-risk genes with known effects are shown along with their risk scores from the SFARI database (1, high confidence; 2, strong candidate; 3, suggestive evidence; and S, syndromic). (**B, C**) Expression patterns for some ASD-risk genes that are specific for (**B**) L30 and (**C**) W30 relative to the non-fighting fish (mean ± SD); light blue color indicates W30 ASD-specific genes and coral color indicates L30 ASD-specific genes. *FDR < 0.05, **\*\***FDR < 0.03; FDR, false discovery rate.

Finally, we performed a cis-regulatory network analysis to test the hypothesis that shifts in neurogenomic states in the W30 and L30 fish were modulated by specific cis-regulatory elements (61). We searched for common transcription factor−binding sites (motifs) in the upstream region corresponding to 1 kb, 3 kb, and 5 kb from the transcription start site in the DEGs obtained from the comparisons B vs. W30 (FDR < 0.05) and B vs. L30 (FDR < 0.05) using HOMER. Most of the significant motifs were found in the 1-kb region, followed by the 3-kb and then 5-kb regions (Table 2).

**Table 2.** Top enriched promoter motifs within the 1-kb, 3-kb, and 5-kb regions upstream of the transcription start site in W30 and L30 DEGs.

| 1-kb |  |  | 3-kb |  |  | 5-kb |  |  |
| --- | --- | --- | --- | --- | --- | --- | --- | --- |
| Matrix | W30 | L30 | Matrix | W30 | L30 | Matrix | W30 | L30 |
|  | p-value | p-value |  | p-value | p-value |  | p-value | p-value |
| HRE | 0.0001 | 0.01 | HRE | 0.01 | NS | HRE | 0.01 | NS |
| GRE | 0.001 | 0.01 | THRb | 0.01 | NS | IRF2 | NS | 0.01 |
| PGR | 0.001 | 0.01 | RORg | 0.01 | NS |  |  |  |
| GATA | 0.01 | 0.001 | GATA | NS | 0.01 |  |  |  |
| HSF6 | 0.01 | NS | BACH1 | NS | 0.01 |  |  |  |
| PRDM14 | 0.01 | NS |  |  |  |  |  |  |
| HSF7 | 0.01 | NS |  |  |  |  |  |  |
| ARE | NS | 0.01 |  |  |  |  |  |  |

HRE: hypoxia-response element; GRE: glucocorticoid response element; IRF2: interferon regulatory factor 2; NS: no significant difference.

Several motifs were enriched and were shared between the W30 and L30 fish, such as the hypoxia-response element (HRE), glucocorticoid response element (GRE), and interferon regulatory factor 2 (IRF2). HRE motifs are used to respond to oxygen stress (hypoxia), and they are recognized by hypoxia-inducible factors (encoded by e*gln* and *hif* genes) (62)—a family of transcription enhancers that play a critical role in the response of the cell to oxygen deficiency (63). Similarly, GRE motifs have a vital role in the cellular response to stress (64). A recent study showed that glucocorticoids promote HIF activity via multiple routes including glucocorticoid receptors (65). Collectively, it seems that both winners and losers were tolerant of oxygen deficiency and stress based on the enrichment of HRE and GRE transcription factor−binding motifs among the DEGs that were expressed in the brains of these fish. Identifying transcription factors predicted to bind to these HRE and GRE motifs using chromatin immunoprecipitation−sequencing is an interesting topic for future studies.

In conclusion, our study has provided insight into the transcriptional profiles of male *B. splendens* brains across different stages of aggressive confrontation, especially with respect to the emergence of a basal neurogenomic state after fighting. In this state, the pair specificity of the brain transcriptome during fighting was gradually lost after the conflict was resolved. This unique neurogenomic state has now been described for the first time from the perspective of the transcriptome and could lay out a framework for exploring the genetic basis of aggression in vertebrates.

## Methods

### Ethics statement

The animal experimentation procedures used in this study were approved by the Institutional Animal Care and Use Committee (IACUC) (Approval No.106171) of the National Cheng Kung University, Tainan, Taiwan.

### Animals and behavioral experiments

Fish collection and experimental procedures have been described previously (26). Briefly, several males of *B. splendens* (average standard length, 5.2 ± 1.1 cm) were imported from a local fish shop in Thailand. When brought to the laboratory for testing, all experimental males were isolated for at least 1 week. All fish were fed with commercial food daily and kept on a 12-hour light/12-hour dark cycle.

For the behavioral trials, each fighting pair, with individual fish distinguished by their colors, i.e., dark red vs. dark blue, was allowed to fight in a 1.7-L PVC tank (18 × 12.5 × 7.5 cm). Five groups of fish were analyzed: (i) non-fighting fish (n = 5 individuals; B1, B2, B3, B4, B5) were used as the control group and were not exposed to other fish; (ii) fish that were allowed to fight for 20 min (D20; n = 5 pairs; D_11 vs. D_12, D_21 vs. D_22, D_31 vs. D_32, D_41 vs. D_42, and D_51 vs. D_52); (iii) fish that were allowed to fight for 60 min (D60; n = 5 pairs; D60_11 vs. D60_12, D60_21 vs. D60_22, D60_31 vs. D60_32, D60_41 vs. D60_42, and D60_51 vs. D60_52); (iv) fish that were allowed to fight just until one fish chased the other (A0; n = 3 pairs; W0_1 vs. L0_1, W0_2 vs. L0_2, and W0_3 vs. L0_3), which usually takes >1 h; and (v) fish that were allowed to fight until one fish chased the other and then were collected 30 min later (A30; n = 3 pairs: W30_1 vs. L30_1, W30_2 vs. L30_2, and W30_3 vs. L30_3) (S1 Fig). We referred to the paired A0 fish as winners 0 (W0) and losers 0 (L0) and to the paired A30 fish as winners 30 (W30) and losers 30 (L30).

For the behavioral analysis, all behavioral experiments were videotaped with a Nikon Cool Pix E5400. Activities were recorded with respect to both the frequency and timing of biting/striking and surface-breathing. Seven fighting pairs in which winners chased losers were analyzed in detail using the video editing software Windows Movie Maker (Microsoft). To examine the aggressiveness levels of the fighting opponents, biting/striking attacks and surface-breathing events were recorded in terms of their frequency and duration. Then the number of biting/striking attacks and surface-breathing events was counted within a 60-min fight. A paired t-test was used to test for the significance of the aggressiveness levels between winners and losers using the *t-test* function in R. A Pearson correlation test was applied to test for a correlation between biting/striking attacks and surface-breathing events across all fighting pairs using the *ggpubr* R package. The figures showing the differences in aggressiveness between winners and losers and the correlation between biting/striking attacks and surface-breathing events were generated using the *boxplot* function in R.

### RNA sequencing (RNA-seq)

For tissue preparation, males for RNA-seq were collected before fighting (B), during fighting (D20 and D60), and after fighting (A0 and A30) (26). Their heads were flash frozen in liquid nitrogen, and their whole brains were carefully dissected and placed individually in Eppendorf tubes containing 1 mL of TRIzol Reagent (Life Technologies). Total RNA was isolated using TRIzol Reagent according to the manufacturer’s recommendation and was subsequently purified on columns with Quick-RNA MiniPrep (Zymo Research, USA). RNA was eluted in a total volume of 30 μL in RNase-free water. Samples were treated with DNase (QIAGEN) to remove genomic DNA. RNA quantity was assessed using a Qubit (Eugene, Oregon, USA), and RNA quality was assessed using the Agilent Bioanalyzer 2100 Nano kit (Agilent, USA) (RNA integrity number RIN: 6.3–8.8). RNA was immediately stored at −80°C until it was used to prepare the sequencing libraries.

RNA-seq libraries were constructed using the TruSeq Stranded mRNA Library Prep kit (Illumina, USA) with proper quality controls, and the molar concentrations were normalized using a KAPA Library Quantification kit (Kapa Biosystems, USA) as described (26). Libraries were sequenced on the Illumina HiSeq 2500 system at Yourgene Bioscience Co., Ltd. (Taipei, Taiwan) and on the Illumina HiSeq 2500 system at the NGS High Throughput Genomics Core (Biodiversity Research Center, Academia Sinica, Taiwan).

### RNA-seq informatics

FASTQC was used to assess the quality of the reads (66). Adaptor sequences and low-quality bases were clipped from 50-bp single-end and paired-end sequences using the Cutadapt tool (67). RNA-seq produced an average of ~28 million reads per sample. We aligned reads to the *B. splendens* reference genome (68) using TopHat version 2.1.1 (69) and Bowtie2 version 2.1.0 (70) with the default settings. The unique mapping reads (reads that matched the reference genome at only one position) were extracted using Samtools (71). The exon-mapped reads were counted with featureCounts (72). The normalized expression levels of genes, represented by the trimmed mean of M-values (TMM), were generated with the edgeR (73) package in R.

To define DEGs, we included genes with at least one count per million (cpm) in at least one sample. Count data were normalized by the TMM (74) using edgeR in R. To assess differential expression, a nested interaction model was fitted in edgeR. A tagwise dispersion estimate was used after computing common and trended dispersions. We adjusted the p-values from all contrasts at once with respect to the false discovery rate (FDR). Two criteria were used to call DEGs: (i) the relaxed version used FDR < 0.05 alone, and (ii) a stringent version used both the FDR and FC value, with FDR < 0.05 and |logFC| > 2; these were implemented by the edgeR package in R.

### Calculation and analysis of CV values

A CV value was calculated for each gene by dividing the standard deviation of its expression by its average expression in each respective group—B, D20, D60, A0, and A30. An ANOVA test followed by post hoc Tukey’s test was used to test for differences in CV values between groups. We used TMM values from all data set**s**(23,411 genes) for all fighting groups (B, D20, D60, A0, and A30) to calculate the CV values.

### Clustering

To differentiate neurotranscriptomic responses between fighting groups, a principal component analysis was conducted using the MASS package (75) in R. To cluster specific DEGs from each fighting group, a heatmap was created using the pheatmap package (76) in R. To visualize brain-transcriptomic synchronization, a GoM model was generated using the CountClust package in R as described (77). Box plots were generated using the *boxplot* function in R. To obtain the overlapping genes between fighting stages, a Venn diagram was constructed using webtools (http://bioinformatics.psb.ugent.be/webtools/Venn/).

A WGCNA was used to find clusters of co-expressed genes (78). Briefly, the weighted average expression profile for each gene module was summarized in an eigengene. Correlations between the eigengene for each gene module and the social phenotypes (B, D20, D60, A0, A30, W0, L0, W30, and L30) and observed behavioral traits were computed to assess the involvement of each module in each social phenotype/behavior. Behavioral traits were classified as aggressive traits (biting, striking, chasing), which were associated with the D20, D60, W0, and W30 fish, or submissive traits (freezing and fleeing), which were associated with the L0 and L30 fish. Genes in each module were subsequently tested for gene ontology using the Database for Annotation, Visualization and Integrated Discovery (DAVID) as described below.

### Statistical analysis

To test whether there was a greater overlap of genes between fighting stages than expected by chance alone, we used a hypergeometric test (23,411 total genes analyzed), which was conducted with the *phyper* function in the dplyr package in R. To obtain the correlation of gene expression values (r), Pearson correlation coefficients of log_2_-transformed TMM values were calculated between individuals from each fighting group (S2 Fig and S3 Fig). A total of 15 pairwise comparisons obtained from the combination of all possibilities between two random individuals among the six individuals of the A0 group or the A30 group using the *pair* function in R. To assess how similar the transcriptomes of paired fish were and to what extent the paired fish were synchronized, we compared the r values for gene expression from paired fish to the r values from unpaired fish from the same treatment (e.g., A0-paired fish vs. A0-unpaired fish; A30-paired fish vs. A30-unpaired fish) using a permutation test, which was implemented with the exactRankTests package in R with all default settings.

### Identifying synchronized genes

The procedure to identify synchronized genes was described in our previous study (26). Briefly, the synchronized genes were characterized by the following two-step method. In the first step, the DEGs from the B vs. A0 and B vs. A30 comparisons (FDR < 0.05) were obtained as described above. In the second step, we divided the procedure into two part as follows. First, to determine whether a particular gene (i) is synchronized or not, the expression distance for each particular gene (Di) between two random fish among the total of 15 possible combinations among which 3 pairs were actual fighting pairs and 12 pairs were randomly paired fish.

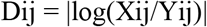

where Dij is the distance for gene i between a random pair j (X and Y), and Xij and Yij are the TMM values for gene i of fish X and fish Y from the pair j, respectively.

Then, a permutation test for each Di compared between the three paired fish vs. the 12 unpaired fish was separately conducted for each of the fighting groups, A0 (A0 paired fish vs. A0 unpaired fish) and A30 (A30 paired fish vs. A30 unpaired fish), to assess the statistical significance. A gene that has a value of p < 0.05 was considered indicative of a synchronized gene. The tests were implemented by the *exactRankTests* package in R with all default settings.

### Identifying learning- and memory-related genes

To find genes related to “memory” or “learning” terms, the sequences from the 2,082 DEGs in L0 fish, 2,843 DEGs in W0 fish, 1,470 DEGs in L30 fish, and 1,150 DEGs in W30 fish relative to the B fish (FDR < 0.05) were searched against the amino acid sequences related to “memory” or “learning” obtained from the UniProt database (79) using blastx with a cut-off E-value of 1e-20. For biological interpretation of synchronized genes, DAVID was used to carry out a Kyoto Encyclopedia of Genes and Genomes (KEGG) pathway analysis (p < 0.05).

### Identifying ASD-risk genes

To identify ASD-risk genes, we first obtained DEGs by comparing the non-fighting group with the L30 and with the W30 groups (FDR < 0.05). This analysis generated 1,470 DEGs and 1,150 DEGs for L30 and W30, respectively. Next, we queried 1,079 ASD-risk genes listed in the Simons Foundation Autism Research Initiative (SFARI) database (sfari.org) (54) in June 2019. Then, we looked for overlap between the L30 DEGs and W30 DEGs and the ASD-risk genes to identify W30-associated ASD-risk genes and L30-associated ASD-risk genes. The potential risk of each gene is represented by its SFARI score, with “1” indicating high confidence of involvement in ASD, “2” indicating a strong candidate for involvement in ASD, “3” indicating suggestive evidence for involvement in ASD, and “S” indicating “syndromic” with respect to autism (see https://gene.sfari.org/ for more detail).

### Gene ontology (GO) analysis

The significantly enriched GO terms (biological process and molecular function terms) and KEGG pathways were identified by DAVID (80). We tested for the overrepresentation of transcripts with a raw p-value of <0.05 (Bayesian statistic). The REVIGO tool (81) was used to filter out GOs obtained from DAVID that were redundant. KEGG mapper tool (82) was used to generate pathways.

### Promoter region analysis

An enrichment analysis of transcription binding sites (TBSs) was carried out using HOMER (83) with a Linux pipeline (available on request). Genomic annotation files were obtained from Beijing Genomics Institute (GBI)28 and were used to define the promoter region for DEGs (FDR < 0.05) in the W30 and L30 groups corresponding to 1, 3, and 5 kb upstream from their transcription start site. A TBS is considered to be significantly enriched if p < 0.05.

### In situ hybridization

The *c-fos* sequence characterized in our previous study was used as template (26). To generate the probe for in situ hybridization, we first designed primers to amplify the *c-fos* sequence (5’ TTCTCCTTCCGTGGACAATC 3’ and 3’ GGGGCTCAAAGTCATGGTTA 5’). The *c*-*fos* probe length is 924 bp. PCR amplification was carried out for 30 s at 94 °C, 30 s at 55 °C, and 40 s at 72 °C for 30 cycles. The PCR fragment was cloned into pGEM-T vector (Promega) and used for the synthesis of cRNA probe using T3 and T7 RNA polymerases (Stratagene) and a digoxigenin labeling mix (Roche, Switzerland).

The preparation of these brains is independent from the process described in the RNA-seq section. Whole *B. splendens* brains collected after fighting for 20 minutes were dissected out, and fixed in 4 % paraformaldehyde (PFA) in phosphate-buffered saline (PBS) overnight at 4 °C. After fixation, these brains were cryoprotected in 20 % sucrose in PBS, embedded in O.C.T. compound (Sakura Finetek), and coronally sectioned at 30 µm using a cryostat (Leica), and sections were then mounted on slides. Whole-brain sections were hybridized with the digoxigenin-labeled cRNA probes at 60°C overnight. After being rinsed with 50% formamide in 2× SSC, the sections were incubated with RNaseA solution (0.2 mg/mL; Invitrogen) at 37°C for 30 min. The sections were then rinsed with 2× SSC and 0.2× SSC and treated with alkaline phosphatase– conjugated anti-digoxigenin (anti-DIG, 1:500; Roche) for 2 h at room temperature. After the sections were washed, positive signals were visualized with NBT-BCIP (Roche) as the chromogenic substrate.

## Acknowledgments

We thank colleagues at National Cheng Kung University (NCKU) including C-L Huang, H-V Wang, S-F Tzeng, H-J Huang, T-Y Chiang and H-S Sun for giving us the opportunity to pursue this study and for useful discussions. We thank Riki Kawamura and Mitsuto Aihara for helping us prepare reagents for wet-lab experiments. This was supported by NCKU, Taiwan with support from the Aim for the Top University Project of NCKU (D104-38A05 & D105-38A03) as well as by the Ministry of Science and Technology in Taiwan (NSC 102-2621-B-006-002- and MOST 103-2621-B-006-005-) to N.O. Computational resources were provided by the Data Integration and Analysis Facility, National Institute for Basic Biology, Japan.

## Supporting information captions

**S1 Video**. Male *B. splendens* fish shift their social status.

**S1 Fig. Experimental design**. Thirty-seven samples were collected at different fighting durations. The samples included non-fighting (B), during fighting (D20 and D60), and after fighting (A0 and A30) fish. W0, winners collected at A0; L0, losers collected at A0; W30, winners collected at A30; L30, losers collected at A30

**S2 Fig. Correlation coefficient values (r) of gene expression in paired fish and unpaired fish of the D20 and D60 groups.** Data from the paired fish are marked in red.

**S3 Fig. Correlation coefficient values (r) of gene expression in paired fish and unpaired fish of the A0 and A30 groups and in un-paired fish from the B group**. The paired fish are marked in red.

S4 Fig. The expression of 289 “geared fighting genes” found in the D20 and A0 fighting groups.

**S5 Fig. Correlation of gene expression changes of the shared DEGs between fighting groups**. Adjacent fighting stages (D20 vs. D60, D60 vs. A0, and A0 vs. A30) are highlighted in orange.

**S6 Fig. Persistent genes found in all fighting stages.** (**A, B**) Two hundred twenty-seven up-regulated genes (**A**) and seven down-regulated genes (**B**) that are persistent across all fighting stages (FDR < 0.05 and |logFC| > 2). These genes are indicated by the red circles.

**S7 Fig. Gene modules significantly associated with winners and losers.** Up-regulated and down-regulated genes are shown in red and green, respectively (TMM ± SE). The corresponding enriched biological process of each gene module is shown in the tables on the right.

**S8 Fig. Overlapping learning and memory genes between winners and losers**. Twenty-nine genes were found in all phenotypes (W0, L0, W30, and L30), which are marked by the red circle; “mem” is the abbreviation for memory.

**S9 Fig. Genes associated with learning and memory in winners and losers.** Up-regulated genes (red) and down-regulated genes (green) are shown. The pathway that triggers learning and memory is marked by a red circle. This pathway was generated using 357 learning and memory genes that were characterized as DEGs in the L0 group.

**S1 Table**. List of DEGs obtained from each fighting stage relative to B (FDR < 0.05). (FC, fold change; CPM, counts per million; FDR, false discovery rate).

**S2 Table**. Comparison of logFC in DEGs between fighting groups.

**S3 Table**. List of biological process terms for up- and down-regulated genes in fighting groups relative to the non-fighting group.

**S4 Table**. List of 289 “geared fighting genes”.

**S5 Table**. The duration and frequency of coloration changes of two fighting opponents within a fighting pair in the initial 20-min fight.

**S6 Table**. The significantly associated gene modules found for each phenotype (B, D20, D60, winners, and losers) as well as for the behavioral traits (aggression and submission).

**S7 Table**. Top 100 genes of gene cluster 3 obtained from GoM analysis.

**S8 Table**. Statistical analysis of the correlation of variation (CV) values.

**S9 Table**. List of synchronized genes in A0 and A30 paired fish.

**S10 Table**. List of overlapping torpor-related genes between mice and *B. splendens*.

**S11 Table**. Learning and memory genes that were enriched in the winners and losers.

